# Positive and negative effects of *CaMCA1* deletion on stress sensitivity and virulence

**DOI:** 10.64898/2026.09.23.753419

**Authors:** Derek Wilkinson

**Author notes:** Corresponding author: Derek Wilkinson.

## Abstract

Metacaspases are cysteine proteases, found in every group of organisms except metazoa. They are structural orthologs of caspases, which orchestrate apoptosis in metazoa. Like caspases (**C**ysteine-dependent **ASP**artate-directed prote**ASES**), paracaspases, and orthocaspases, metacaspases are class C14 proteases, with a hemoglobinase fold and catalytic cysteine residue. However, metacaspases cleave after arginine or lysine instead of aspartate, leading some to call for metacaspases to be renamed. Furthermore, most metacaspases are activated by calcium, they do not target the same range of proteins as caspases, and they tend to cut target proteins multiple times rather than at a single, canonical site. The *Candida albicans* metacaspase, Mca1p mediates cell death in response to various stresses but also possesses pro-life functions such as clearing protein aggregates and lifespan extension. Many plant and protist metacaspases lack cell death roles and mediate development, differentiation and immunity. This article highlights a reversed effect of Mca1p on growth inhibition by acetic acid, hydrogen peroxide and amphotericin B and on virulence of *C. albicans*, when injected into *Galleria mellonella* (wax moth) larvae, depending on whether cells originate from exponential or stationary phase culture. This reversal of metacaspase function hints at the protein’s dual nature and at the conditions that drive the switch from its pro-survival to its pro-death role.

## 1. Introduction

Caspases orchestrate regulated cell death (RCD), development, differentiation, inflammation and positive and negative selection of immune cells in metazoa^[1,2]^. Distant orthologs of caspases were identified in animals, plants, fungi, archaea, and bacteria ^[3-6]^. These orthologs are called paracaspases, metacaspases and orthocaspases, and researchers debated whether metacaspases were true functional orthologs of caspases ^[7]^. It is now known that they have different substrate specificities, usually require calcium for activation of proteolysis and that many have only pro-survival functions, though several appear to have pro-survival and pro-death functions ^[8]^.

At the turn of the century, Madeo and coworkers showed that *S. cerevisiae* undergoes regulated cell death in response to mutations in the cell cycle machinery and certain stresses, that this cell death resembles metazoan apoptosis and involves new protein biosynthesis and that hydrogen peroxide-induced cell death is dependent on the metacaspase Mca1p/Yca1p ^[9,10]^.

Phillips et al. ^[11]^ demonstrated that 5-10 mM hydrogen peroxide (H_2_O_2_), 40-60 mM acetic acid (HAc) or 4-8 μg/mL amphotericin B (AmB) induced a form of cell death in *C. albicans* that resembled apoptosis. Cao et al. ^[12]^ showed that deleting *MCA1* in *C. albicans* reduced sensitivity to cell death caused by oxidative stress, reduced intracellular ROS accumulation, increased trehalose accumulation, lowered ATP levels and lowered mitochondrial membrane potential. They suggested that more sugar phosphates were being fed into trehalose synthesis rather than oxidative phosphorylation, leading to reduced ATP synthesis and reduced reactive oxygen species (ROS) production by mitochondria.

The identification of regulated cell death and orthologs of the metazoan cell death machinery in *C. albicans* is exciting, as candidemia is one of the top four hospital-acquired bloodstream infections; *C. albicans* is the main agent of candidemia, and mortality among candidemia patients may be over 40% ^[13,14]^. Targeting components of the fungal cell death machinery could lead to the development of novel classes of antifungal drugs ^[11]^.

*Galleria mellonella* (wax moth) larvae are widely accepted as an alternative model for testing fungal virulence and the efficacy of antifungal drugs ^[15]^since it avoids the ethical problems associated with mammalian models, is cheaper and larval upkeep is very simple, with larvae eating filter paper in the bottom of a Petri dish. Fungal suspension may be injected into the pro-leg, and experiments take only one or two days. Several varieties of hemocytes resemble neutrophils, and virulence results resemble those in BalbC mice. However, larvae lack human-like organs, so they cannot be used as models of invasion of the heart, kidneys, spleen etc.

In this study, it is shown that deleting the *C. albicans* metacaspase gene *MCA1* reduces the rate of stress-induced cell death and increases the virulence of *C. albicans*, injected into *Galleria mellonella* (wax moth) larvae. This is consistent with reports in the literature that metacaspases reduce stress resistance and virulence by promoting stress-induced cell death ^[5,16]^. However, when cells in these experiments were derived from stationary culture, the deletion of *MCA1* did not affect stress-induced cell death but decreased virulence, which is consistent with reports of a pro-survival role ^[8]^. Possible reasons for this phenomenon are discussed.

## 2. Materials and methods

Media components were from Formedium (Swaffham, Norfolk, UK) and other reagents from Fisher Scientific (Pittsburgh, Pennsylvania, USA) unless stated otherwise.

### 2.1 Strains, plasmids, culture and storage

*Candida albicans* strains used in this study are shown in Table 1 and include strain SN78 ^[17]^ as the parental background, as well as three prototrophic strains, produced in this study: the *MCA1* wild type (WT), the *mca1Δ*/*mca1Δ* double deletion mutant (Mut) and the *MCA1* reintegrant strain (Reint), with *MCA1* integrated into the *RPS1* locus of the Mut background. *Escherichia coli* strains used in this study for plasmid production were mainly in the DH5α background ^[18]^, and are shown in Table 2. The CIp40 plasmid was transformed into Strataclone SoloPack competent *E. coli* cells (Agilent Technologies, Santa Clara, California, USA).

**Table 1.**
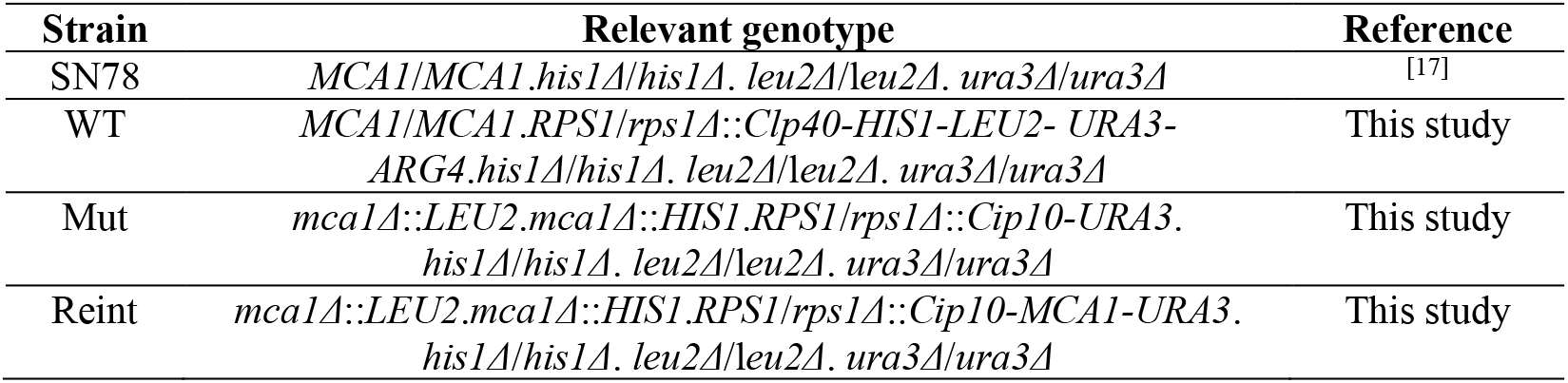
C. albicans strains used in this study.

| Strain | Relevant genotype | Reference |
| --- | --- | --- |
| SN78 | *MCA1/MCA1.his1Δ/his1Δ. leu2Δ/leu2Δ. ura3Δ/ura3Δ* | [17] |
| WT | *MCA1/MCA1.RPS1/rps1Δ::Clp40-HIS1-LEU2- URA3- ARG4.his1Δ/his1Δ. leu2Δ/leu2Δ. ura3Δ/ura3Δ* | This study |
| Mut | *mca1Δ::LEU2.mca1Δ::HIS1.RPS1/rps1Δ::Cip10-URA3. his1Δ/his1Δ. leu2Δ/leu2Δ. ura3Δ/ura3Δ* | This study |
| Reint | *mca1Δ::LEU2.mca1Δ::HIS1.RPS1/rps1Δ::Cip10-MCA1-URA3. his1Δ/his1Δ. leu2Δ/leu2Δ. ura3Δ/ura3Δ* | This study |

**Table 2.** Plasmids used in this study.

| Plasmid | Auxotrophy marker | Resistance marker | Reference |
| --- | --- | --- | --- |
| pSN52 | *HIS1* | KanR<sup>1</sup> | [17] |
| pSN40 | *LEU2* | KanR<sup>1</sup> | [17] |
| Cip10 | *URA3* | AmpR<sup>2</sup> | [19] |
| Clp30 | *HIS1-URA3-ARG4* | AmpR<sup>2</sup> | [20] |
| Clp40 | *HIS1-LEU2-URA3-ARG4* | AmpR<sup>2</sup> | This study |
| Cip10-*MCA1* | *URA3* | AmpR<sup>2</sup> | This study |

<sup>1</sup> Kanamycin resistance.
<sup>2</sup> Ampicillin resistance.

*C. albicans* strains were grown in liquid culture in YPD (1% (w/v) yeast extract, 2% (w/v) peptone, 2% (w/v) D-glucose) and an aliquot of culture was mixed with an equal volume of 50% (v/v) glycerol/water for long term storage at -80 °C. When needed, frozen stock was streaked onto YPD agar (YPD + 2% (w/v) agar) using a sterile pipette tip and incubated at 30 °C for two days. Overnight cultures were prepared by adding one colony from the agar plate to 10 mL YPD using a sterile pipette tip and incubating at 30 °C overnight. To produce exponential (mid-log phase) cells for transformations and for virulence or stress sensitivity testing, 50 mL fresh YPD was inoculated with 1 mL overnight culture and incubated at 30 °C for 4 hours. To produce stationary phase cells, the 50 mL liquid culture was incubated at 30 °C for 2 days. For selective culture, 200 µL cell suspension was spread onto selective agar (6.9 g/L yeast nitrogen base with ammonium sulphate but without amino acids; 2 % (w/v) D-glucose; 2 % (w/v) agar No. 2; 0.72 g/L amino acid dropout mix without leucine, histidine, or uridine; 50 mg/L leucine and/or histidine and/or uridine as appropriate). *Escherichia coli* was grown in lysogeny broth (LB: 1% (w/v) tryptone, 0.5% (w/v) yeast extract, 1% (w/v) sodium chloride) and aliquots were mixed with equal volumes of 50% (v/v) glycerol/water and stored at -80 °C until needed. *E. coli* was streaked onto LB agar (LB + 2% (w/v) agar) and incubated at 37 °C overnight, and 10 mL LB was inoculated with a colony taken from the agar plate and incubated at 37 °C. Where required, kanamycin or ampicillin was added to LB or LB agar at a final concentration of 50 and 100 µg/mL, respectively.

### *2*.*2 C. albicans* transformation

*Candida albicans* cells were transformed with cassettes made *via* polymerase chain reaction (PCR) amplification of auxotrophy markers on plasmids ^[17]^ or with linearized plasmids containing auxotrophy markers ^[19,20]^. The transformation protocol was adapted from that of Gietz and Woods ^[21]^. Cells from a 50 mL mid-log phase culture (above) were washed with 50 mL LATE (10 mM Lithium Acetate, 10 mM Tris, 1mM EDTA pH 8.0 in sterile milliQ water) and resuspended in 1 mL LATE, then 100 µL aliquots were incubated overnight with 0.7 mL PLATE (40% (w/v) PEG (polyethylene glycol) in LATE), 80 µL DNA and 5 µL boiled herring sperm (10 mg/mL). The next day, cells were heat-shocked for one hour at 42 °C, then cells were washed in water, resuspended in 400 µL of water, spread on two selective agar plates, and incubated at 30 °C for 2-3 days, until colonies appeared.

### 2.3. E.coli transformation

50 µL of frozen competent cells were defrosted on ice, 5 µL of chilled DNA suspension was added and the tube was tapped to encourage mixing. The tube was left on ice for half an hour then cells were heat shocked at 42 °C for 45 seconds and placed on ice for a further 2 minutes. 1 mL of LB, preheated to 42 °C, was added to the cells; the cell suspension was incubated for 60 mins at 37 °C with shaking (200 rpm). Two aliquots (50 µL and 500 µL) were spread on LB agar (LB + 2% agar) plates containing 100 µg/mL ampicillin or 50 µg/mL kanamycin, as appropriate, and incubated overnight at 37 °C.

### 2.4. Miniprep

Plasmid harvesting from *E. coli* was conducted using a miniprep kit (Quiagen, Crawley, West Sussex, UK) in accordance with the manufacturer’s instructions.

### 2.5. Gel electrophoresis

An electrophoresis gel (0.7% agarose and 0.5 µg/mL ethidium bromide in TAE buffer (40 mM Tris acetate, 1 mM EDTA, pH 8.5)) and DNA was pipetted into the wells. Then a current of 100 V was applied for 20 to 30 minutes. A Gbox gel imager (Syngene, Cambridge, UK) was used to visualize DNA bands in the gel. Images were recorded on thermal print paper and saved to file as appropriate.

### 2.6. Gel extraction

Under a UV light the appropriate DNA band was cut out of the gel with a scalpel and the gel slice placed in a clean Eppendorf tube. Gel purification was conducted with a gel extraction kit (Qiagen, Crawley, West Sussex, UK) in accordance with the manufacturer’s instructions.

### 2.7 PCR

25 μL of PCR Master Mix (Thermo Scientific, Waltham, Massachusetts, USA) (final concentration: 0.625 units Taq DNA polymerase, 75 mM Tris-HCl [pH 8.8], 20 mM ammonium sulphate, 1.5 mM magnesium chloride, 0.01% Tween ® 20, 0.2 mM of each dNTP), 22 μL of sterile nuclease-free water, 1 μL of forward primer (0.2 μM), 1 μL of reverse primer (0.2 μM) (Table 3) and 1 μL (0.5 to 125 ng) of DNA template (Table 2) were mixed in a PCR tube and polymerase chain amplification carried out in a Thermal Hybaid thermal cycler (Franklin, Massachusetts, USA) using the following cycle: initial denaturation at 95 °C for 5 minutes; 30 cycles of i) denaturation at 95 °C for 30 seconds, ii) annealing at 53 °C for 1 minute and iii) extension at 72 °C for 3 minutes; followed by a final extension at 72 °C for 5 minutes.

**Table 3.**
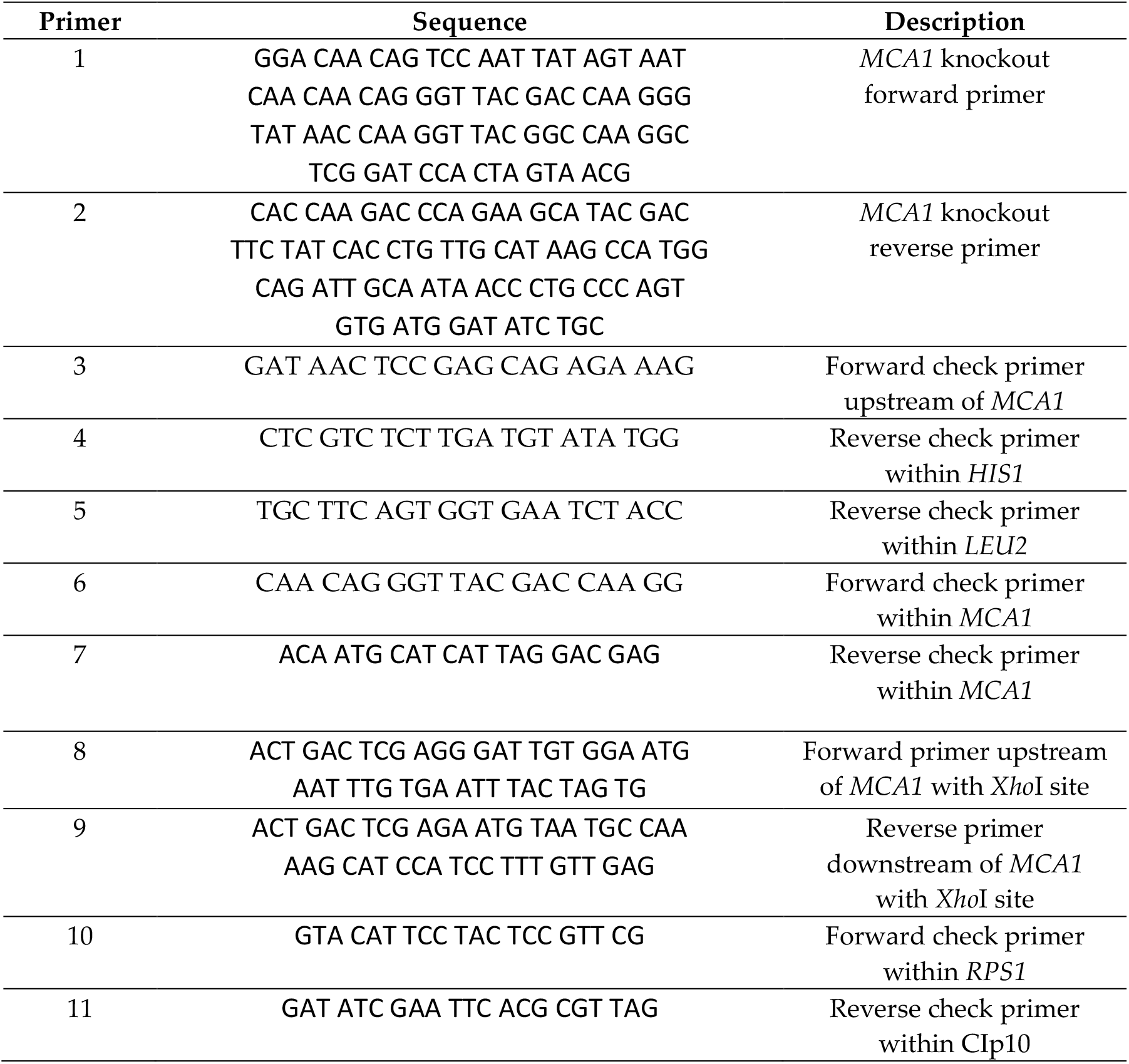
PCR primers used in this study.

### 2.8. Colony PCR

A single colony was resuspended in 25 µL of sterile milliQ water. 5 µL of cell suspension was streaked on fresh agar (LB or YPD, as appropriate) and incubated at 37 °C overnight (*E. coli*) or 2 days at 30 °C (*C. albicans*) to prepare stock for freezing. Meanwhile, the same *E. coli* cell suspension was used directly as DNA for PCR amplification, but the *C. albicans* cell wall had to be digested first. 10 mg/mL of lyticase (Thermo Scientific, Waltham, Massachusetts, USA) was added to the tube of *C. albicans* cell suspension and incubated at 37 °C for 10 minutes, followed by freezing at -80 °C for 5 minutes and thawing at room temperature. The treated suspension was then applied to PCR amplification.

### 2.9. Strain and plasmid creation

The *Candida albicans* parental strain used in this study was SN78, developed by Noble and Johnson ^[17]^, which is derived from *C. albicans* clinical isolate SC5314 and was deleted for both copies of the *HIS1, LEU2* and *URA3* genes and was therefore auxotrophic for histidine, leucine and uridine (Table 1). Noble and Johnson also developed plasmids to be used in gene knockouts (Table 2: pSN52 and pSN40). Since the plasmids contained the *HIS1* and *LEU2* genes respectively, transformants could be selected on YPD agar plates lacking histidine and/or leucine. Each strain was restored to prototrophy by incorporating the appropriate linearized plasmid (Cip10 ^[19]^, Cip10-MCA1 or Cip40 [Table 2]) into the *RPS1* locus.

#### 2.9.1 Prototrophic wild type (WT)

Plasmid Clp30: ^[20]^ (a kind gift from Dr. Steve Bates) contains the *HIS1, URA3* and *ARG4* genes while *C. albicans* strain SN78 ^[17]^ lacks the *HIS1, LEU2* and *URA3* genes. To restore HIS/LEU/URA prototrophy, a new plasmid, Clp40, containing *HIS1, LEU2, URA3* and *ARG4* was created from Clp30 and integrated into the *RPS1* locus of SN78. First, the *LEU2* gene of plasmid pSN40 was amplified by PCR using primers 1 and 2 (Table 3), the DNA was precipitated and washed. It was digested with *Bsa*BI (New England Biolabs, Ipswich, MA, USA), A-tailed, and purified using a gel extraction kit (NBS Biologicals, Huntingdon, UK) in accordance with the manufacturer’s instructions. DNA was cloned into Strataclone vector pSC-A amp/kan and transformed into Strataclone SoloPack competent *E. coli* cells (Agilent Technologies, Santa Clara, California, USA). The plasmid was checked *via Spe*I digestion and gel electrophoresis. The *LEU2* gene was flanked by two *Spe*I restriction sites. The *Spe*I restriction fragment was ligated into *Xba*I-digested Clp30 to yield Clp40. The Clp40 plasmid was linearized by digestion with *Stu*I and transformed into strain SN78, incorporating the auxotrophy markers into the *RPS1* locus to restore prototrophy and yield strain WT (Table 1). Transformants were selected on agar plates lacking histidine, leucine and uridine and checked *via* PCR with primers 10 and 11 and gel electrophoresis.

#### 2.9.2 Prototrophic double *MCA1* deletion strain (Mut)

The *HIS1* gene of plasmid pSN52 and the *LEU2* gene of plasmid pSN40 (Table 2) were amplified by PCR using primers 1 and 2 (Table 3), with 70 bp of homology with the sequences just upstream and downstream of *MCA1*. Successive rounds of transformation using *MCA1* knockout cassettes, created from pSN52 and pSN40, yielded a strain with both histidine and leucine prototrophy. Transformants were selected on agar plates lacking histidine (1^st^ round) and leucine (2^nd^ round). Transformation was checked by PCR amplification using primer pairs 3/4 and 3/5, respectively, followed by gel electrophoresis. The absence of any *MCA1* gene was confirmed by PCR with primers 6 and 7, followed by gel electrophoresis. CIp10 was linearized with *Stu*I and transformed into the double knockout strain. Transformants were selected on plates lacking histidine, leucine, and uridine. The resulting prototrophic strain was named Mut (Table 1).

#### 2.9.3 Prototrophic *MCA1* reintegrant strain (Reint)

Lyticase treatment was used to obtain SN78 cell contents and 1 µL used for PCR amplification of *MCA1* with primers 8 and 9, yielding *MCA1*, flanked by *Xho*I restriction sites. The *Xho*I digestion fragment was ligated into *Xho*I-digested Cip10. Gel extraction was used to purify the CIp10-*MCA1* plasmid, which was transformed into DH5α competent *E. coli* cells (Invitrogen, Waltham, Massachusetts). Successful transformation was confirmed by *Xho*I digestion and gel electrophoresis. CIp10-*MCA1* was linearized with *Stu*I before transformation into the Mut strain to yield strain Reint (Table 1), with *MCA1* in the *RPS1* locus. Transformants were selected on agar plates lacking uridine and successful integration of CIp- *MCA1* was confirmed by PCR with primers 10 and 11 (a kind gift from Dr. Steve Bates), followed by gel electrophoresis.

#### 2.10 Stress-induced growth inhibition

The WT, Mut and Reint strains were evaluated for sensitivity to regulated cell death (RCD)-inducing levels of hydrogen peroxide (H_2_O_2_), acetic acid (HAc) and amphotericin B (AmB). *C. albicans* cells from mid-log phase (4-hour-old) or stationary phase (48-hour-old) culture were centrifuged, washed in water, and resuspended at 1 x 10^5^ cells/mL in water. 1 mL of cell suspension was spread on a YPD agar plate and dried. A sterile 5 mm filter paper disc was placed in the center of each plate. 5 μL of 7.5 M HAc, 2.5 M H_2_O_2_ or 2 mg/mL AmB was pipetted onto the disc, and the plate was incubated at 30 °C for 1 day. The diameter of the zone of fungal growth inhibition around the disc was measured, and the area was calculated as diameter squared multiplied by 3.14. Three biological replicates (cells from independent cultures, derived from different colonies). The area of the zone of inhibition (ZOI) for the Mut strain was compared to those of the WT and Reint strains and the ZOIs of the WT and Reint strains were also compared. ANOVA analysis and Turkey testing were carried out using Jasp version 0.98.1 ^[22]^. Since three strains and three pairwise comparisons were included in each analysis, the Bonferroni correction was applied to a confidence interval of 95%, and the threshold for significance was adjusted from 0.05 to 0.0167.

#### 2.11 Virulence in wax moth larvae

Cells from mid-log phase (four-hour old) or stationary phase (48-hour old) YPD liquid cultures were washed with phosphate-buffered saline (PBS: 136.89 mM sodium chloride; 2.68 mM potassium chloride; 10.14 mM dibasic sodium phosphate; 1.8 mM monobasic potassium phosphate; pH 7.4) and resuspended in PBS at 3 x 10^7^ cells/mL. 10 µL of cell suspension or 10 µL of PBS was injected into the left, front proleg of each wax moth larva. Larvae were placed in a Petri dish, lined with filter paper, and incubated at 37 °C. Live and dead larvae were counted every 6 hours. Ten larvae, chosen at random, were injected with each strain (WT, Mut or Reint) and ten with PBS. Live (pale and moving) and dead (melanized and unmoving) larvae in each plate were counted, and the numbers were recorded before checking the strain label on the bottom of the plate. The experiment was carried out three times on separate occasions with independent yeast cultures and different batches of larvae. Kaplan-Meier curves were generated and log-rank testing carried out in R (version 4.6.1 ^[23]^) using packages survival version 3.8.11 ^[24]^ and survminer version 2.5.2 ^[25]^.

#### 2.12 Effect of Mca1p on serum-induced filamentation

Cells from stationary (48 hour-old) cultures of WT, Mut and Reint were centrifuged, washed in water, and resuspended at a density of 1 x 10^7^ cells/mL in YPD + 10% (v/v) fetal calf serum (FCS). Cell cultures were incubated with shaking (200 rpm) at 37 °C, and samples were examined at regular intervals beneath a light microscope. For each strain, 400 cells in 4 fields of vision were examined, and the numbers of yeast and hyphae were counted. The experiment was conducted three times on different occasions. Means and standard deviations were calculated in Excel and plotted as bar graphs and student’s t-tests were carried out, comparing the Mut with WT and with Reint and comparing WT with Reint.

#### 2.13 Effect of Mca1p on growth rate

Cells from mid-log phase cultures of the WT, Mut and Reint strains were washed and resuspended in synthetic complete medium (6.9 g/L yeast nitrogen base with ammonium sulphate and amino acids; 2 % (w/v) D-glucose; 2 % (w/v) agar No. 2) to achieve an absorbance at 650 nm (A650) of 0.2. Then 10 µL of SC pH3 was added to each well of a microtiter plate and media in successive columns included 240 mM, 120 mM, 60 mM, 30 mM, 15 mM and 0 mM acetic acid. 100 µL of cell culture was added to each well. The final HAc concentrations were 120 mM, 60 mM, 30 mM, 7.5 mM, and 0 mM. The plate was placed in a VersaMax automatic plate reader with Softmax Pro 5.4.1 software (Molecular Devices, Sunnyvale,\ California, USA) and incubated at 30 °C for 24 hours. Absorbance at 650 nm was read and recorded every 3 minutes. Raw data was exported to and processed in Excel. Mean absorbance for each triplicate reading was calculated and normalized to the mean absorbance at time zero. Absorbances and natural logs of absorbances were plotted against time. The linear portions of semi-logarithmic curves were used to calculate the doubling time (generation time, G) using equation (I).

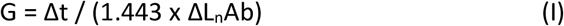

where Δt = time passed and ΔL_n_Ab = change in natural log of absorbance Due to time constraints, the experiment was only carried out once.

#### 2.14 Effect of Mca1p on acetic acid-induced cell death in yeast and hyphae

Cells from stationary phase cultures of WT, Mut and Reint were centrifuged, washed and resuspended at 1 x 10^7^ cells/mL in SC pH3 + 10% (v/v) FCS and either 80 mM HAc or an equal volume of water. After incubation for 3 hours at 37 °C with shaking (200 rpm), cells were centrifuged, washed and resuspended in 10 µg/mL propidium iodide in PBS and incubated at 37 °C for half an hour. Cells were washed and resuspended in PBS then examined using a fluorescence microscope (Leica DMLB, Wetzlar, Hesse, Germany) with an excitation energy of 488 nm and a band pass filter of 562-588 nm. Dead (red PI-stained) and live (unstained) cells were counted. Treatment with FCS at 37 °C induced germination, and mother (yeast) cells had attached daughter hyphae. The numbers of live and dead mother and daughter cells were counted.

## 3. Results

### 3.1. Mca1p mediates stress-dependent growth inhibition in *C. albicans* cells from mid-log phase culture

The wild type (WT), *mca1*Δ/*mca1*Δ double knockout mutant (Mut) and *MCA1* reintegrant (Reint) strains were spread on agar plates and dried. Filter paper discs were placed in the centers of the plates and treated with RCD-inducing concentrations of H_2_O_2_, HAc or AmB. Three biological replicates (cells from independent cultures, derived from separate colonies) were included for each strain. After incubation at 30 °C for 24 hours, the diameter of the zone of inhibition (area of no fungal growth) around each disc was measured. The means and standard deviations of three independent experiments are shown (Figure 1A, C, E). ANOVA analyses and Turkey tests were conducted for each treatment/culture phase comparison to compare results for the Mut strain with those for WT and Reint. There were significant differences (p < 0.0167) between WT and Mut and between Reint and Mut but not between WT and Reint (p > 0.0167) regarding sensitivity to HAc (Figure 1A), H_2_O_2_ (Figure 1C) and AmB (Figure 1E) when spread cells were derived from mid-log phase culture. In each case, the *MCA1* double knockout strain Mut was less sensitive to the cell death-inducing stressor than either of the strains (WT and Reint) with functional *MCA1* genes.

**Figure 1.**
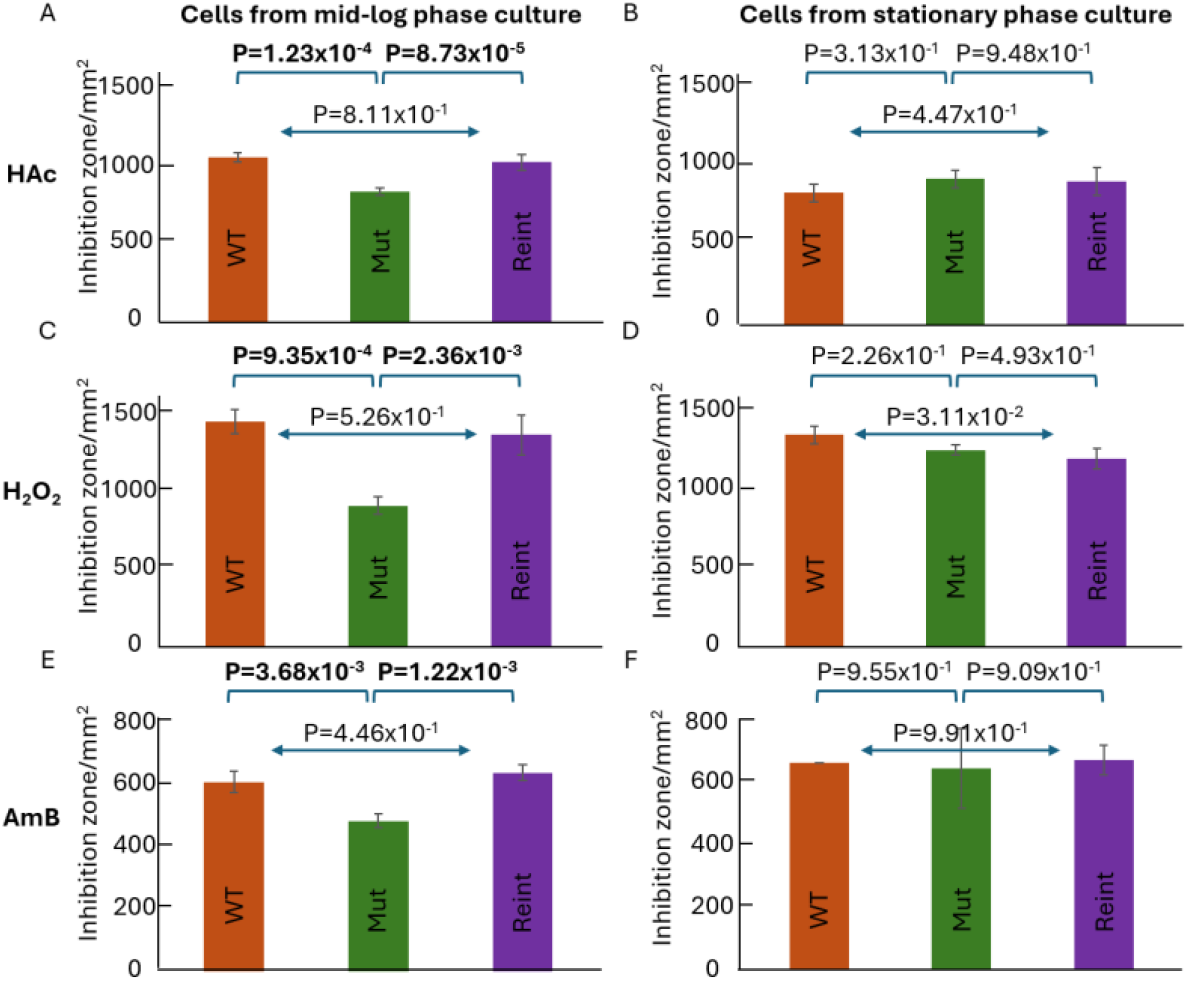
Role of Mca1p in growth inhibition by HAc, H_2_O_2_ and AmB. (A) Cells from mid-log phase culture treated with HAc. (C) Cells from mid-log phase culture treated with H_2_O_2_. (E) Cells from mid-log phase culture treated with AmB. (B) Cells from stationary-phase culture treated with HAc. (D) Cells from stationary phase culture treated with H_2_O_2_. (E) Cells from stationary phase culture treated with AmB. *C. albicans* wild type (WT: brown), double *MCA1* knockout (Mut: green) or *MCA1* reintegrant (Reint: purple) strains were spread on agar and dried. A 5 mm diameter filter paper disc was placed in the center of each plate and treated with 5 µL of 7.5 M HAc (A and B), 2.5 M H_2_O_2_ (C and D) or 2 mg/mL AmB (E and F). After 24 h incubation at 30 °C the diameter of the zone of inhibition (ZOI) around each disc was measured and the area calculated. Three biological replicates were used for each strain, and the means (bars) and standard deviations (error bars) are shown above. Anova analysis and Turkey testing were carried out for each treatment/culture phase combination. The pairwise comparison to-which each p value refers is indicated by a bracket or double-headed arrow above the bar chart. Bold: significantly different (p < 0.0167).

### 3.2. Mca1p does not mediate stress-dependent growth inhibition in *C. albicans* cells from stationary phase culture

The cell death induction assay was repeated with cells from stationary phase (48 hours old) cultures (Figure 1B, D and F). There were no significant differences (p > 0.0167) among the mutant (Mut), wild type (WT), and reintegrant (Reint) strains regarding sensitivity to HAc (Figure 1B), H_2_O_2_ (Figure 1D), and AmB (Figure 1F) when spread cells were derived from stationary phase culture.

### 3.3. Mca1p reduces the virulence of mid-log phase-derived *C. albicans* cells

To investigate the role of Mca1p in *C. albicans* virulence, cells from log-phase cultures of WT, Mut and Reint were washed and resuspended in PBS. Cell suspensions (or PBS alone in the control) were injected into wax moth larvae (ten per strain in each experiment), and the larvae incubated at 37 °C. The numbers of live and dead larvae were counted blind at 6 hourly intervals. The experiment was carried out three times with independent yeast cultures (i.e. three biological replicates). Kaplan-Meier curves were generated and log-rank testing carried out in R (version 4.6.1 ^[26]^) using packages survival version 3.8.11 ^[24]^ and survminer version 2.5.2 ^[25]^. The curves and p-values from statistical comparisons of pairs of curves are shown in Figure 2A. Generally, Mut killed larvae more rapidly than WT or Reint, (p = 7 x 10^- 7^ and p = 9 x 10^-5^, respectively), while there was no statistical difference between survival curves for the WT and Reint strains (p = 0.3). None of the control larvae, injected with PBS, died during the 36-hour period. All larvae died within 36 h of inoculation — but larvae died more quickly when injected with *C. albicans* that lacked Mca1p.

**Figure 2.**
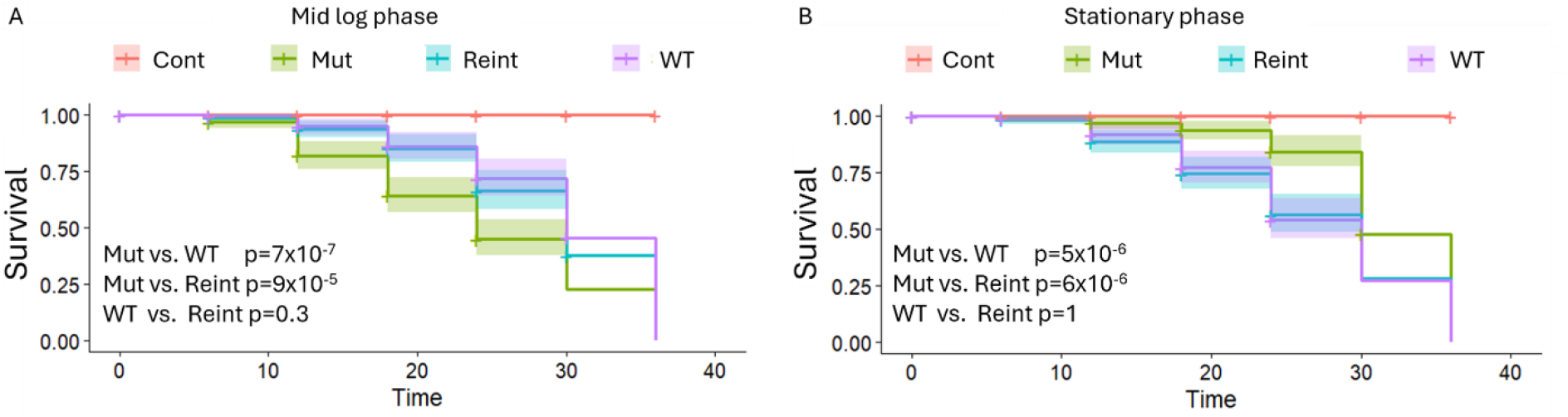
Role of Mca1p in virulence. (A) *C. albicans* cells from mid-log phase culture. (B) *C. albicans* cells from stationary phase culture. Strains: wild type (WT: lilac line), double *MCA1* knockout (Mut: green line) or *MCA1* reintegrant (Reint: blue line). Cells were suspended in PBS and injected into wax moth larvae and incubated at 37 °C. Control larvae were injected with PBS alone (Cont: orange line). Larval survival was checked every 6 hours until all *C. albicans*-injected larvae had died. Three independent experiments were conducted. Kaplan-Meier curves were plotted and log-rank testing carried out in R. Lines represent estimated cumulative survival probabilities over time and shading around lines represents 95% confidence interval. p-values: probability that the difference between two curves occurred by chance

### 3.4. Mca1p increases the virulence of stationary phase-derived *C. albicans* cells

The virulence assay was repeated with *C. albicans* cells from stationary phase culture (Figure 2B). Again, all PBS-injected larvae survived for 36 hours after inoculation. In contrast with the previous experiment, Mut appeared to kill larvae more slowly than WT or Reint (p = 5 x 10^-6^ and p = 6 x 10^-6^, respectively), but there was no significant difference between Kaplan-Meier curves for the WT and Reint strains (p = 1). Unexpectedly, injection with Mut increased larval survival, rather than decreasing it. All larvae died within 36 h of inoculation; but larvae died more slowly when injected with *C. albicans* that lacked Mca1p.

### 3.5. Mca1p appears to have no effect on serum-induced filamentation

Mca1p appeared to affect the virulence of *C. albicans* in wax moth larvae. A major virulence factor of *C. albicans* is its ability to switch from yeast to hyphal form and escape from innate immune cells ^[27-29]^. To test whether Mca1p influenced *C. albicans* virulence *via* altered filamentation, cells from stationary-phase cultures of each of the three strains (WT, Mut and Reint) were grown at 37 °C and 200 rpm in YPD plus 10% FCS, and the numbers of yeast and hyphae were counted at 15-30 min intervals. Means and standard deviations were calculated and plotted as bars and error bars (respectively) of bar charts (Figure 3). There was no significant difference (p > 0.01 in student’s t-tests) in the rate of filamentation of any of the strains, compared with any of the others at any time point.

**Figure 3.**
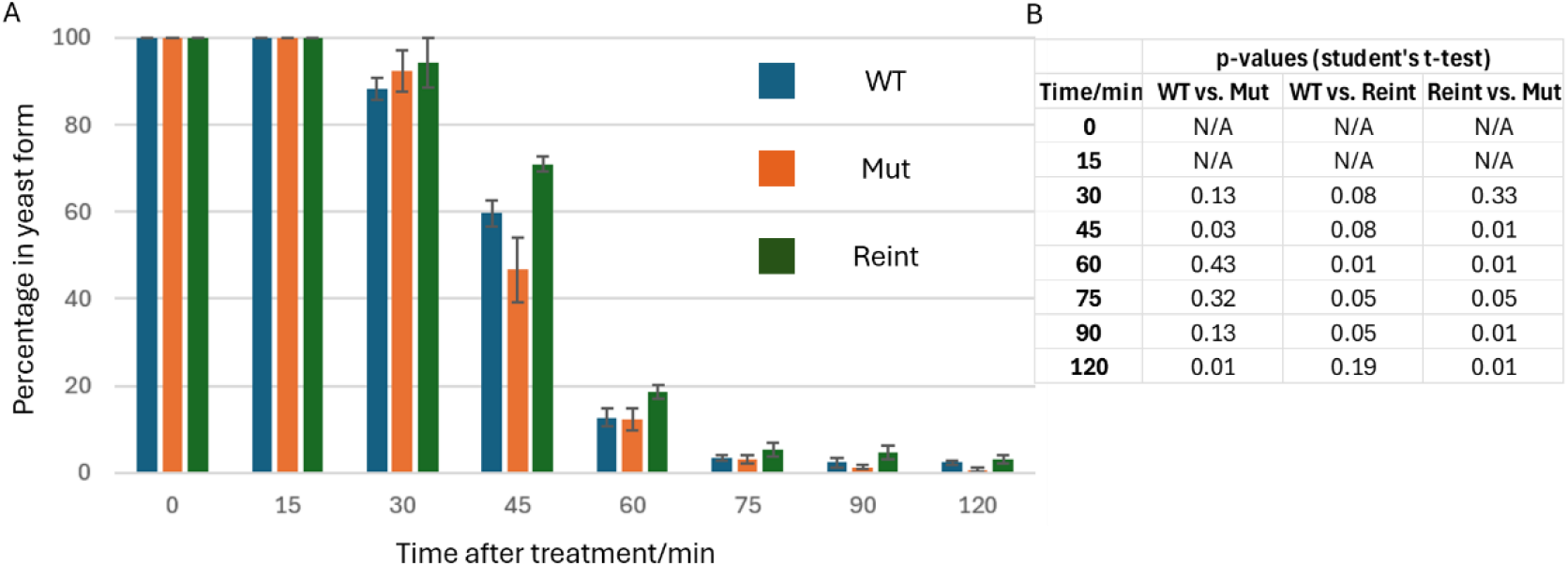
Role of Mca1p in serum-induced filamentation. (A) Percentage of cells in yeast form (rather than hyphal form) at various time points after serum treatment. Three strains were used: Wild type strain (WT: blue); *MCA1* double deletion mutant strain (Mut: orange); *MCA1* reintegrant strain (Reint: green). Cells from stationary-phase culture were grown in YPD + 10% FCS at 37 °C. At 15-30 min intervals, samples were examined beneath a microscope, and the numbers of yeast and hyphae were counted. At least 400 cells in 4 fields of vision were examined for each strain. Three independent experiments were conducted, and the bars and error bars (above) represent mean percentages of cells in yeast, rather than hyphal, form; and standard deviations, respectively. B. p-values from Student’s t-tests, comparing each strain with each other strain at individual time points. No p-value was generated for times 0 and 15 min after treatment since percentages were identical.

### 3.6. Mca1p has no effect on the percentages of yeast or hyphae that die during acetic acid treatment

One of the possible explanations for differences in virulence between cells with and without a functional *MCA1* gene could be different rates of filamentation and different sensitivities in yeast and hyphae to stresses, encountered in the host. Therefore, yeast of all three strains (WT, Mut and Reint) were induced to undergo filamentation by treatment with SC (pH 3) + 10% FCS at 37 °C in the presence or absence of 80 mM acetic acid. Then propidium iodide was used to stain dead cells and a fluorescence microscope used to count live and dead cells (mother yeast and/or daughter hyphae). The results were plotted as bar charts (Figure 4). M+ (live mother cells) and M- (dead mother cells) add up to 1; H+ (live hyphae) plus H- (dead hyphae) also add up to 1. Finally, M+H+ (live mother/live hypha) plus M+H- (live mother/dead hypha) plus M-H+ (dead mother/live hypha) plus M-H- (dead mother/dead hypha) add up to 1. Chi-squared testing was conducted to identify significant differences in percentage cell death among the 3 strains. Cell death was significantly higher among treated than untreated hyphae (p < 0.0001) and among treated than untreated yeast (p < 0.0001). There was no significant difference between Mut and WT cell death among treated mother cells (p = 0.34) or treated hyphae (p = 0.48). However, a significantly greater proportion of Mut hyphae and yeast died than those of WT or Reint (p < 0.0001) when not treated with acetic acid.

**Figure 4.**
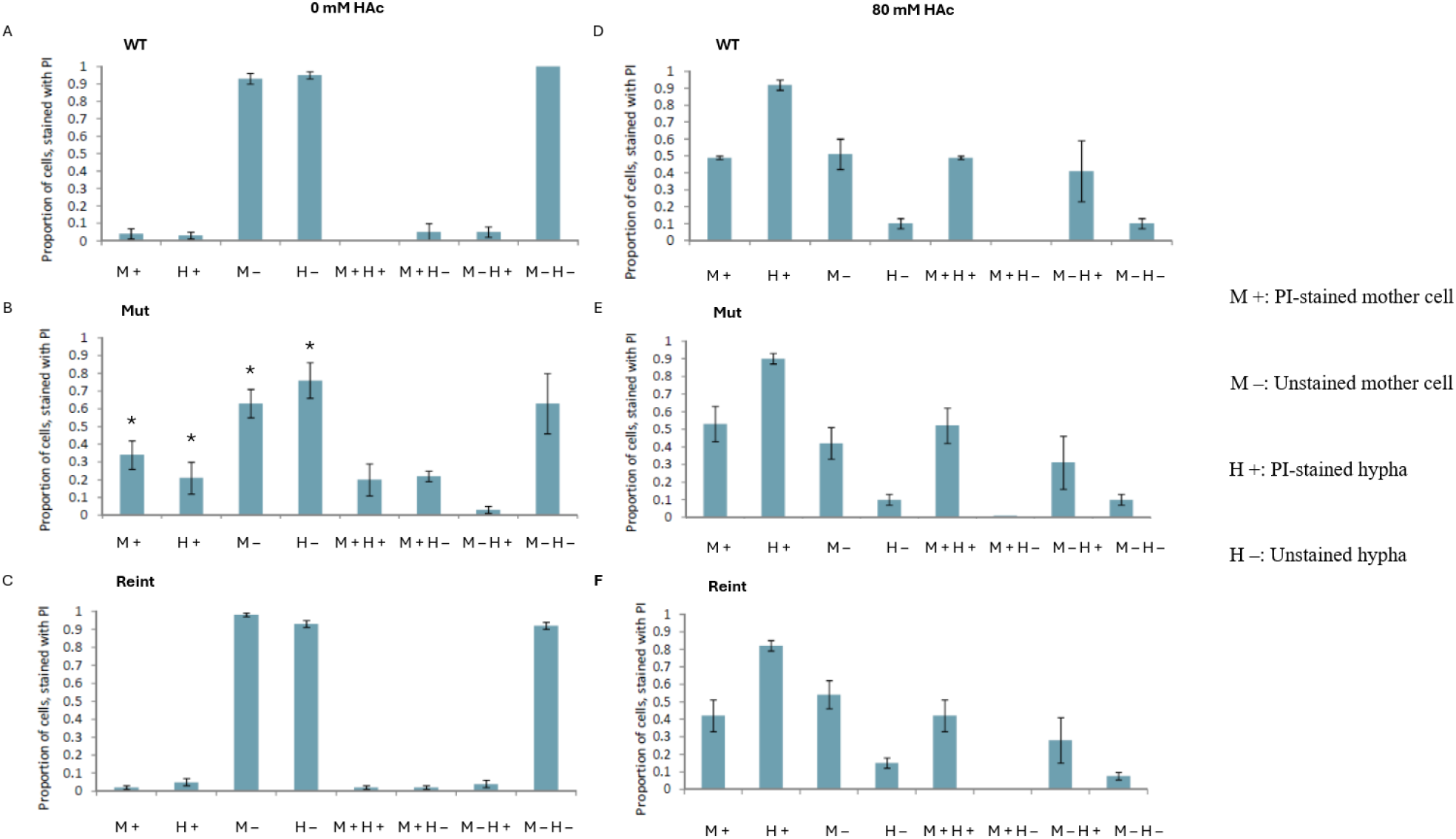
Cell death of yeast and hyphal cells in acetic acid-treated and untreated *Candida albicans*. (A) Untreated WT. (B) Untreated Mut. (C) Untreated Reint. (D) Treated WT. (E) Treated Mut. (F) Treated Reint. Cells from stationary-phase culture were washed and resuspended at 10^7^ cells/mL in synthetic complete (SC) medium (pH 3) with 10% FCS and either 80 mM HAc or an equal volume of water. After 3 h incubation at 37 °C with shaking (200 rpm) the cells were washed in PBS and resuspended in 10 µg/mL propidium iodide and incubated at 37 °C for 30 min. After washing and resuspending in PBS, cells were examined using a fluorescence microscope. Live (non-fluorescent) and dead (red, fluorescent) hyphal daughter and yeast mother cells were counted, as were combinations of stained or unstained mother and stained or unstained daughter cells. M+: stained (dead) mother (yeast) cell; M–: unstained (living) mother (yeast) cell; H+: stained (dead) hyphae (daughter cells); H–: unstained (living) hyphae (daughter cells). Asterisk (*): significant difference (p < 0.0001) between Mut and WT and between Mut and Reint.

## 4. Discussion

Three RCD-inducing stressors (hydrogen peroxide, acetic acid, and amphotericin B) were tested against strains with (WT and Reint) and without (Mut) a functional *MCA1* gene. Mca1p mediated growth inhibition in response to each stressor when cells were derived from log-phase culture (Figure 1A, C and E) but played no such role in cells derived from stationary-phase culture (Figure 1B, D and F). Growth inhibition may involve regulated cell death (RCD), accidental cell death (e.g. necrosis) or negative regulation of the cell cycle. Other studies have demonstrated a role for Mca1p in fungal cell death, induced by hydrogen peroxide, acetic acid and amphotericin B ^[30,31]^. Though not tested here, possible reasons for the absence of this pro-death function in cells from stationary phase, compared with mid-log phase, culture include differences in Mca1p expression, differences in Mca1p proteolytic activity and/or protein aggregate clearance or stress adaptation in stationary phase culture that counteract the pro-death function of Mca1p. It was previously shown that Mca1p expression is highly induced by RCD-inducing concentrations (160 mM) of acetic acid but only modestly induced by lower, stress-inducing (20 mM) concentrations ^[32]^. It is possible that differing conditions within the cell during log-phase and stationary-phase growth lead to differing levels of stress and modulate expression of Mca1p. Murphy et al. ^[33]^ showed that Mca1p abundance did not change appreciably during the transition from log to stationary phase. However, *MCA1*/Mca1p expression was not tested at the mRNA or protein level in this study. Future experiments might include measuring expression in cultures over several time points.

The decreased rate of larval death when injected with strains with a functional *MCA1* gene (WT and Reint) rather than a strain with no *MCA1* gene (Mut) is consistent with a role for Mca1p in the mediation of cell death or growth inhibition, induced by elements of the host immune system (such as ROS, RNS and low pH). In moths and butterflies, granulocytes and plasmatocytes phagocytose smaller microbes and encapsulate larger invaders ^[34]^. The phagolysosomes of both cell types kill microorganisms *via* reactive oxygen and nitrogen species, low pH, antimicrobial peptides, etc., and these hemocytes also secrete ROS and RNS ^[34]^. If Mca1p mediates acid- and peroxide-induced cell death, *C. albicans* cells with a functional *MCA1* gene might be more sensitive to wax moth immune attack than cells with no *MCA1* gene. It has been shown that metacaspases mediate stress-induced cell death in pathogenic protists and reduce virulence in animal hosts ^[10]^. Mechanisms, not tested in this study, by which metacaspases could affect virulence might include altered growth rate, increased sensitivity to immune attack, changes in the rate at which cells switch morphology, and differences in the sensitivity of morphotypes. It should be stressed that no analysis of fungal load or other indicators of pathogenesis in the larvae was carried out. Therefore, it is not possible to comment upon the precise mechanism by which some strains promote larval death at a different rate from others.

Cao et al. ^[12]^ found that deleting *MCA1* resulted in slower growth of *C. albicans* in either rich medium or synthetic medium. The strains used by Cao et al. were derived from strain CaI4, rather than SN78, but both parental strains were derived from clinical strain SC5314. The medium used (SC or SD) was similar. The *MCA1* deletion and reintegration techniques were slightly different but should not affect growth rate. Based on the results of the Cao et al. study, it is possible that virulence differences in wax moth larvae were due to the effect of Mca1p on growth rate.

There was no significant difference in the rate of serum-induced switching from yeast to hyphal form in the three strains (Figure 3), so Mca1p appears to play no role in this process and morphology changes may not be the reason behind altered virulence. In acetic acid-treated hyphae or yeast (Figure 4), there were no significant differences in the percentages of cells that died. However, among untreated cells, hyphal and yeast cell death was higher in the mutant (Mut) strain than the wild type (WT) or reintegrant (Reint) strains. It should be stressed that the bar charts in Figure 4 represent the proportions of mother cells (yeast) or daughter cells (hyphae) that were alive or dead and the proportion of mother/daughter pairs that were live/live, live/dead, dead/live, or dead/dead. Since propidium iodide staining was used to identify dead cells, only those with a compromised plasma membrane were detected. Any dead cells with intact membranes would be counted as live in this experiment. Mca1p has been shown previously to extend lifespan in aging yeast cells ^[35]^. The effects of hydrogen peroxide and other ROS on cell death among hyphae or yeast were not tested in this study, so it cannot be ruled out that differences in the virulence of strains with and without a functional *MCA1* gene were due to differences in either hyphal or yeast sensitivity to host hemocyte ROS, RNS, low pH, etc. However, this does not explain the increased virulence when cells were derived from stationary-phase culture. It was shown previously that *Candida albicans* hyphae were more resistant than yeast cells to AmB and that this phenomenon was dependent on metacaspase Mca1p ^[36]^. It cannot be ruled out that a specific stressor, present in wax moth larvae, affects fungal growth in a metacaspase-dependent manner.

While the loss of a pro-death function in cells derived from stationary-phase, rather than mid-log-phase, culture is interesting (Figure 1), the reversal of Mca1p’s role in wax moth larvae from an antagonist to an agonist of virulence when fungal cells were derived from stationary phase, rather than mid-log phase, culture (Figure 2) is remarkable. Future experiments could test the effect of culture phase on the cleavage of known Mca1p targets.Experiments have shown that fungi can adapt to stressful conditions, such as increased acetic acid concentration, that this adaptation includes upregulated expression of proteins such as catalase and that adaptation protects against stress-induced RCD ^[37]^

It should be stressed that *MCA1* is incorporated into the *RPS1* locus rather than the native locus in the reintegrant and that this may affect the rate of expression. Throughout the experiments in this study, the reintegrant and the wild-type behaved differently from the mutant strain but similarly to each other. However, differences in expression (and therefore possibly in behavior) cannot be ruled out. In future experiments, *MCA1* could be integrated into the native locus, ruling out this source of uncertainty.

## 5. Conclusion and future prospects

Mca1p mediates acetic acid-, hydrogen peroxide-, and amphotericin B-induced growth inhibition in *C. albicans*, but this effect is absent in *C. albicans* cells from stationary culture. Mca1p also reduces virulence of *C. albicans* in wax moth larvae, possibly *via* increased sensitivity to elements of the host immune system. This phenomenon reverses when *C. albicans* cells are derived from stationary-phase culture, which is consistent with previous reports of dual pro-death and pro-survival roles, though this has not been directly tested in this study.

More research needs to be conducted to establish why Mca1p appears to switch behavior in mid-log phase and stationary phase cells. A comparison of the relative abundances of Mca1p in log phase and stationary phase cultures and the proportions of Mca1p molecules that are cleaved and/or localized to protein aggregates would also be useful. The rate of killing by mammalian phagocytes of cells from log phase and stationary phase culture and testing of cell death among yeast and hyphae under different stresses should be assayed using cells from log phase and stationary phase culture. The effects of stress adaptation on metacaspases could be further researched as this could explain the loss of Mca1p function in stress-induced RCD. Another useful experiment would be to measure intracellular free calcium in log-phase and stationary-phase *C. albicans* cells. Finally, screening a gene knockout library for sensitivity to cell death in log-phase and stationary-phase *C. albicans* cells could help to identify the pathways that regulate cell death and the precise role of Mca1p and therefore potential targets for antifungal drugs. Metacaspase inhibitors have been successfully used to kill bloodstream forms of Plasmodium and Trypanosoma species ^[38,39]^. However, such inhibitors target pro-survival or differentiation-promoting metacaspases. Fungal metacaspases may have pro-survival and pro-death roles, and any drug would have to promote the pro-death role or block pro-survival activity. At least one drug (the lipopeptide, C17 Fengycin B, shows promise against *Fusarium oxysporum* by activating metacaspase-dependent cell death ^[40]^. Other potential targets include the calcium efflux system in the endoplasmic reticulum or calmodulin, since calcium promotes the cell death activity of some metacaspases ^[8]^. However, more research is needed to elucidate the signaling pathways that regulate the different activities of fungal metacaspases.

